# Transcriptome analyses of mouse cardiac myocytes and non-cardiomyocytes: postmitotic vs. proliferative cells

**DOI:** 10.1101/2023.08.20.554007

**Authors:** Yasuhiro Takenaka, Masataka Hirasaki, Ikuo Inoue, Masaaki Ikeda, Hisayuki Ohata, Yoshihiko Kakinuma

## Abstract

Adult heart mostly contains long-lived postmitotic cardiomyocytes and non-cardiomyocytes that have proliferative potential. Here, we isolated cardiomyocytes and non-cardiomyocytes from young and aged mouse heart, and performed transcriptome analyses by RNA sequencing to understand the differences of gene expression in postmitotic and proliferative cells. Gene ontology analyses revealed that genes associated with inflammatory response were upregulated in aged cardiac myocytes, whereas genes including two ATP synthases in mitochondrial respiratory complex V (*Atp5l* and *Atp5J2*) and two NADH dehydrogenases in complex I (*Ndufa11* and *Ndufv3*) were significantly downregulated. In aged non-cardiomyocytes, genes related to inflammatory responses were also upregulated, while genes involved in cell cycle and DNA replication process were downregulated. We also found that the expression levels of some small nucleolar RNAs (snoRNAs) are decreased cardiomyocytes with aging. snoRNAs are deeply involved in RNA modification such as pseudouridylation stabilizing ribosomal RNA (rRNA) and mRNA splicing. Therefore, the age-related reduction in snoRNA expression may lead to the destabilization of rRNA, splicing dysfunction, and ultimately a decrease in protein synthesis capacity. A comparison with transcriptome results obtained for non-cardiomyocytes suggests that the decline in the expression of mitochondria-related genes and snoRNAs accompanying aging is specific to cardiomyocytes, implying their potential utility as one of novel aging markers in postmitotic cells.

## Introduction

Cellular senescence is basically defined as the limited replicative potential of normal diploid cells, which is so-called the “Hayflick limit”. The irreversible cell cycle arrest observed in cultured cells is one of the most essential hallmarks of cellular senescence, which is presently considered to be caused by shortened telomeres that trigger a DNA damage response (DDR). Cellular senescence is also characterized by morphological changes such as a hypertrophic cellular shape, increased levels of senescence-associated secretory phenotype (SASP), enhanced activity of the mTOR pathway [1], and expression of senescence-associated β-galactosidase (SA-β-gal) [2]. Therefore, studies of cellular senescence have been predominantly attempted to elucidate the molecular mechanism of irreversible cell cycle arrest and senescence-related phenomenon in proliferating cells. Cellular senescence is fundamentally contributing to organismal aging. Selectively eliminating senescent cells by senolytic drugs restores multiple age-associated disorders such as cardiovascular dysfunctions in aged animals [3, 4]. Senolytic drugs that cause cell death of senescent cells exhibiting proliferation-related markers such as p16^Ink4a^ and p21 [4]. However, many tissues in mammals such as brain, retina, and bone are largely composed of postmitotic terminally differentiated cells and limited number of proliferative cells.

Recent reports suggest that postmitotic cells also show phenotypic characteristics of cellular senescence, and play important roles in organismal aging or contribute to pathology [5]. Adult heart mostly contains postmitotic/infrequently dividing cardiac myocyte and cardiac non-myocyte cells including endothelial cells, fibroblasts, erythrocytes and leukocytes that have proliferative potential [6]. Several studies have shown that senescent phenotypes are observed in cardiomyocytes and other cell types with aging or under age-related pathological condition [7–9] although roles and contribution of senescence in each cell type to heart degeneration and cardiac disease are not well-defined [10, 11]. The analyses of gene expression profiles of aged cardiomyocytes have been performed to elucidate the overall physiological aging and aging-related diseases of the heart [7, 12, 13]. However, there are very few studies that directly compare the gene expression profiles between rodent cardiomyocytes and non-cardiomyocytes with aging. Here, we isolated cardiomyocytes and non-cardiomyocytes from young and aged mouse heart, and performed transcriptome analyses by RNA sequencing to understand the differences of gene expression in postmitotic and proliferative cells.

## Materials and methods

### Animals

The experimental protocols in this study were approved by the Animal Care and Research Ethics Committee of Nippon Medical School in Tokyo, Japan (permission number: 2022-006). All animal experiments were performed in strict accordance with the recommendations set by the ARRIVE guidelines and carried out in accordance with National Institutes of Health guide for the care and use of laboratory animals. Animal stress and suffering were minimized as much as possible. Male C57BL//6J mice were purchased from Tokyo Laboratory Animals Science Co., Ltd. Mice were housed in a pathogen-free condition and 12 h light and dark cycles with ad libitum access to standard chow diet (CREA Japan).

### Preparation of cardiac myocyte and non-cardiomyocytes suspension from mouse heart

Mouse cardiomyocytes and non-cardiomyocytes were prepared by the method based on Ackers-Johnson *et al.* [14] and Kosloski *et al.* [15] with slight modifications. After anesthetizing mouse with avertin, abdominal aorta was cut, the heart was exposed, and perfused by injection of 7 mL Perfusion Buffer [Hank’s balanced salt solution (HBSS-) containing 10 mM taurine, 10 mM 2,3-butanedione monoxime (BDM), 2 mM ethylenediaminetetraacetic acid and 0.5 mM α-tocopherol] into the right ventricle using peristaltic pump P-1 (Pharmacia biotech) at 1 mL/min. Ascending aorta was clamped with Reynolds forceps, and the heart was transferred to a 60-mm culture dish containing HBSS. The heart was further injected with 10 mL Perfusion Buffer into the left ventricle using P-1 at 1 mL/min. Enzyme digestion was performed by sequential injection of 22 mL Digestion Buffer [HBSS(-) containing 10 mM taurine, 10 mM BDM, 1.26 mM CaCl_2_, 0.5 mM MgCl_2_, 0.4 mM MgSO_4_, 0.1 mg/mL dispase II (Fujifilm Wako Chemicals), 1.0 mg/mL collagenase 2, 1.0 mg/mL collagenase 4 (Worthington), and 0.5 mM α-tocopherol] using P-1 at 1 mL/min at ambient temperature. Tissue was then transferred into 60 mm-dish of 3 mL fresh Digestion Buffer, minced into ∼1 mm^3^ pieces using forceps, and triturated by gentle pipetting. Cellular suspension was filtrated through a 100-µm strainer, mixed with 5 mL ice-cold Stop Buffer (Perfusion Buffer containing 5% fetal bovine serum), and centrifuged at 13 × g for 1 min at 4°C. The pellet and the supernatant were saved as the cardiac myocyte and non-cardiomyocyte fractions, respectively. The pelleted cells of the cardiac myocyte fraction were resuspended in ice-cold 7 mL Perfusion Buffer and applied onto a 20-µm nylon net membrane (Millipore, SCNY00020). Trapped cells on the membrane were collected with 7 mL Perfusion Buffer, centrifuged at 150 × g for 5 min at 4°C, and pelleted cells were saved as the cardiomyocytes. The supernatant of non-cardiomyocyte fraction was also passed through a 20-µm nylon net membrane. The filtrate was centrifuged at 150 × g for 5 min at 4°C and pelleted cells were saved as the non-cardiomyocytes.

### Cell culture

Isolated cardiomyocytes and non-cardiomyocytes were cultured in in M119 (Sigma-Aldrich) supplemented with 1 mg/mL bovine serum albumin (Sigma-Aldrich), 1×ITS-G supplement (Fujifilm Wako Chemicals), 10 mM BDM, and 1×CD lipid concentrate (Thermo Fisher Scientific), and penicillin-streptomycin (Nacalai Tesque) in a laminin-precoated 24-well plate at 37°C in a humidified atmosphere of 5% CO_2_ based on the Ackers-Johnson’s method [14].

### Flow cytometry (FCM)

Isolated non-cardiomyocytes were analyzed as previously described by Stellato *et al.* [16]. Non-cardiomyocytes resuspended in Cell Staining Buffer (phosphate-buffered saline containing 2% FBS and 0.1% sodium azide) were treated with anti-mouse CD16/CD32 antibody (BioLegend, #101302) for 30 min on ice, followed by incubation with the antibodies against TER119-PE (BioLegend, #116207), CD45-PE (BioLegend, #103105), CD31-PE (BioLegend, #102407), and Podoplamin (gp38)-APC (BioLegend, #127409) diluted in Cell Staining Buffer for 30 min on ice. 7-AAD (BioLegend, #420403) was used to gate live cells. For the Ki-67-staining, cells resuspended in Cell Staining Buffer were stained with Zombie NIR (BioLegend, #423105) for 10 min at 25°C, then fixed in 70% ethanol at −30°C for 2 h. Pelleted cells were washed with Cell Staining Buffer twice, resuspended in Cell Staining Buffer, and stained with FITC-conjugated mouse anti-Ki-67 antibody (BD Biosciences, #556026) for 30 min at 25°C. Cells were washed, resuspended in an appropriate volume of Cell Staining Buffer, and analyzed using the BD FACS Canto II (BD Bioscience). Obtained data was analyzed using FlowJo ver. 10 (BD Bioscience).

### RNA extraction and quantitative PCR (qPCR)

Total RNA was isolated from pelleted cardiac myocyte and non-cardiomyocytes using ISOGEN reagent (Nippon gene) according to the manufacturer’s instructions. The yield and quality of the purified RNA were evaluated by spectrophotometry, agarose gel electrophoresis, and Bioanalyzer assessments. In the RNA sequencing experiment, total RNA was extracted from the cardiac myocyte and non-cardiomyocytes of three mice, and 1 µg of RNA from each mouse was pooled and used for the analysis. For the qPCR experiment, reverse transcription was performed using total RNA extracted from individual mouse cardiac myocyte and non-cardiomyocytes. Briefly, 0.5 µg of total RNA was polyadenylated by poly(A) polymerase (New England Biolab) and reverse-transcribed using ReverTra Ace qPCR RT Master Mix with gDNA Remover (TOYOBO). The cDNA of each mouse gene was amplified and detected using THUNDERBIRD Next SYBR qPCR Mix (TOYOBO) and Thermal Cycler Dice Real Time System III (TAKARA BIO, Shiga, Japan). 18S ribosomal RNA was used for the reference gene. Primer sequences used are available in the Supporting Information.

### RNA sequencing (RNA-seq)

The RNA-seq analysis was conducted using single samples (n=1). The library preparation and RNA-seq were performed by Macrogen Japan (Tokyo, Japan) using a TruSeq^TM^ Stranded Total RNA Library Prep Kit with a Ribo-Zero^TM^ kit and NovaSeq6000 paired-end sequencing (Illumina). The sequencing reads were aggregated into an rRNA reference (Mouse_rRNA_Reference_BK000964) using Bowtie package to remove rRNA reads. Unpaired reads were removed using FASTQ-pair. Paired reads were aligned using STAR version 2.7.8a with the GRCm38 reference alignment. RSEM was used to quantify the transcript. Transcripts Per Million (TPM) calculated using RSEM were log_2_(count+1) transformed.

### Gene Ontology (GO) and Gene set enrichment analysis (GSEA)

To characterize the molecular and functional aspects of the differentially expressed genes (DEGs), GO analyses were performed using the online DAVID database (https://david.ncifcrf.gov/). Using TMM values, which are normalized count data, GSEA was performed according to the methods described on the GSEA website (http://www.gsea-msigdb.org/gsea/index.jsp). Dotplot and MA plot figures were generated using R version 4.2.3.

### Statistical analyses

The two groups were compared using the non-parametric Mann-Whitney *U* test. Differences between the groups were considered significant at *p* < 0.05. All data were analyzed using the GraphPad Prism 5.0 software (GraphPad Software, San Diego, California USA, www.graphpad.com) and presented as the mean (SD) of the obtained values.

## Results

### Gene expression profiling of cardiac myocyte and non-cardiomyocytes isolated from young and aged mice

To isolate cardiac myocyte (CM) and non-cardiomyocytes (non-CM) from the whole heart of young and aged mice, we modified previously described isolation methods [14, 15]. The single cell suspension obtained by passing through a 100 µm cell strainer was further divided into cardiac myocyte and non-CM fractions with a 20 µm cell strainer (Fig. 1A). To assess the appropriateness of the obtained fractionations, expression levels of genes characteristic of cardiac myocyte and non-CM were evaluated by qPCR. In both young and aged mice, cardiac myocyte-specific genes (*Tnnc1* and *αMHC*) and cardiac pacemaker cell-specific gene (*Hcn4*) were highly expressed in the cardiac myocytes (CM) fraction, whereas fibroblast-specific (*Vim* and *Col1a1*) and smooth muscle cell (*αSMA*)-specific genes were predominantly expressed in the non-CM fraction (Figs. 1B and 1C). These results suggest that the fractionation of CM and non-CM was successful in the experiment. We next examined proliferative potential of non-CM by two-color flow cytometry using anti-Ki-67 antibody conjugated with FITC. The single cell suspension was first gated in FSC-A and SSC-A plot (P1 in Fig. 1D), then selected by Zombie NIR staining to exclude dead cells (P2), and finally counted as Ki-67-positive (P3) in histogram. To establish the P3 gate, we prepared cells treated with FITC-conjugated normal mouse IgG as a negative control. Subsequently, FCM analysis was performed, and the threshold for the P3 gate was set at the rightmost point on the FITC histogram where the FITC-signal intensity of the cells treated with IgG became nearly 0. Percentages of Ki-67-positive cells against P2-gated cells were significantly higher in young non-CM fraction (Fig. 1E), suggesting the presence of proliferating cells in the non-CM fraction. In fact, part of isolated young non-CM steadily proliferated under cell culture condition (Fig. 1F).

**Figure 1.**
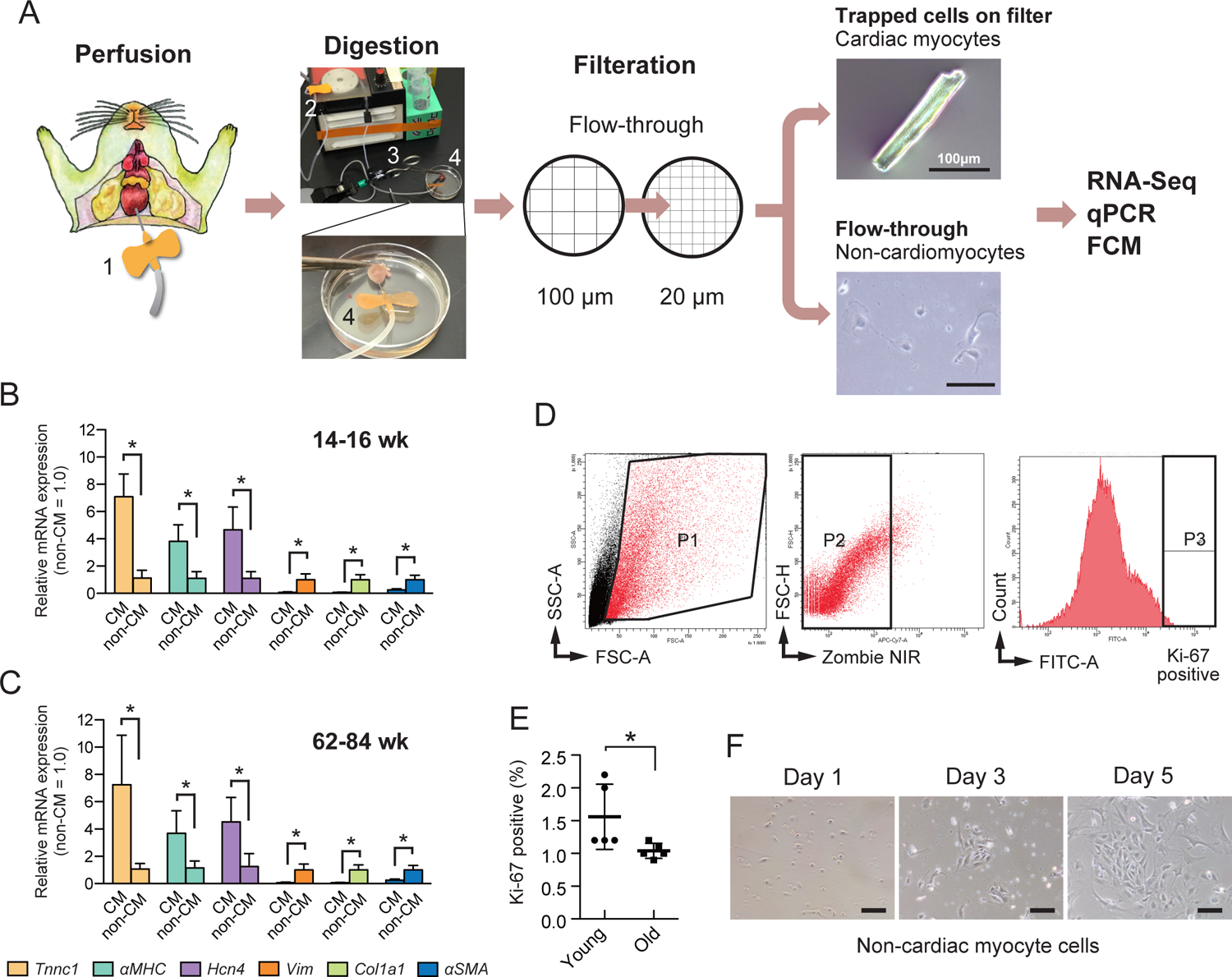
Isolation and evaluation of the cardiac myocytes (CM) and the non-cardiomyocytes (non-CM) fractions from young and aged mice. A, Summary of the isolation protocol. 1, The right ventricle was punctured with a winged needle for perfusion using Perfusion Buffer. The image of dissected mouse upper body is from TogoTV (© 2016 DBCLS TogoTV, CC-BY-4.0 https://creativecommons.org/licenses/by/4.0/deed.ja). 2, Peristaltic pump P-1 used for perfusion. 3, Reynolds forceps to clamp ascending aorta. 4, Clamped heart submerged in 60 mm-dish of Perfusion Buffer. After tissue dissociation with Digestion Buffer, the cell suspension was filtered through a 100 µm-nylon mesh to remove undigested tissue debris. Flow-through-cell suspension was filtered again through a 20 µm-nylon mesh to separate CM (trapped on the filter) and non-CM (flow-through) cells. Collected cell fractions were centrifuged and pelleted cells were used for transcriptome, qPCR, and FCM analyses. Scale bar = 100 µm. Evaluation of the CM and the non-CM fractions isolated from young (B, 14-16 wk) and aged (C, 62-84 wk) mice by qPCR using cardiac myocyte-specific (*Tnnc1* and *αMHC*), cardiac pacemaker cell-specific (*Hcn4*), fibroblast-specific (*Vim* and *Col1a1*) and smooth muscle cell-specific (*αSMA*) primer sets. Relative values are shown as the mean (SD) (*n* = 7). The asterisk (*) indicates *p* < 0.05 by a Mann-Whitney *U* test. D, Representative FCM analysis of proliferative cells in the non-CM fraction based on the Ki-67 marker. P1, the single cell fraction. P2, Zombie NIR-negative live cells. P3, Ki-67-positive cells in histogram. E, Significant reduction in Ki-67-positive cells in the non-CM fraction with aging. Relative values are shown as the mean (SD) (*n* = 5). The asterisk (*) indicates *p* < 0.05 by a Mann-Whitney *U* test. F, Representative images of the non-CM proliferating in culture on days 1, 3, and 5 after isolation from the heart of a young mouse. Scale bar = 100 µm.

### Composition of cardiac fibroblast and non-fibroblast cells were not altered between young and aged non-CM fractions

Next, we investigated whether the proportion of cells comprising the non-CM fraction, such as cardiac fibroblasts and endothelial cells, changes with aging. Type I collagen-producing cardiac fibroblasts are characterized by being negative for Ter119, CD45, and CD31 (lineage negative, Lin^-^) and positive for stromal cell marker gp38 (gp38^+^) [16]. As shown in Fig. 2A, vast majority of cells in non-CM fraction were gp38-negative. The average proportion of gp38^+^ fibroblasts (P3 gate in Fig. 2A) in the aged non-CM fraction (4.8%) was marginally lower than that in the young non-CM fraction (7.0%), although no statistical significance was observed between the two groups (Fig. 2B). We further roughly assessed the proportions of Ter119-(erythrocytes)(Fig. 2C), CD45-(leukocytes)(Fig. 2D), CD31-(endothelial cells)(Fig. 2E) positive, and Lin^+^ (Fig. 2F) cells in the non-CM fractions of both young and aged samples. Our results revealed no significant differences between these fractions, suggesting that the cellular composition of cardiac erythrocytes, leukocytes, endothelial cells, as well as fibroblasts remained largely consistent with aging.

**Figure 2.**
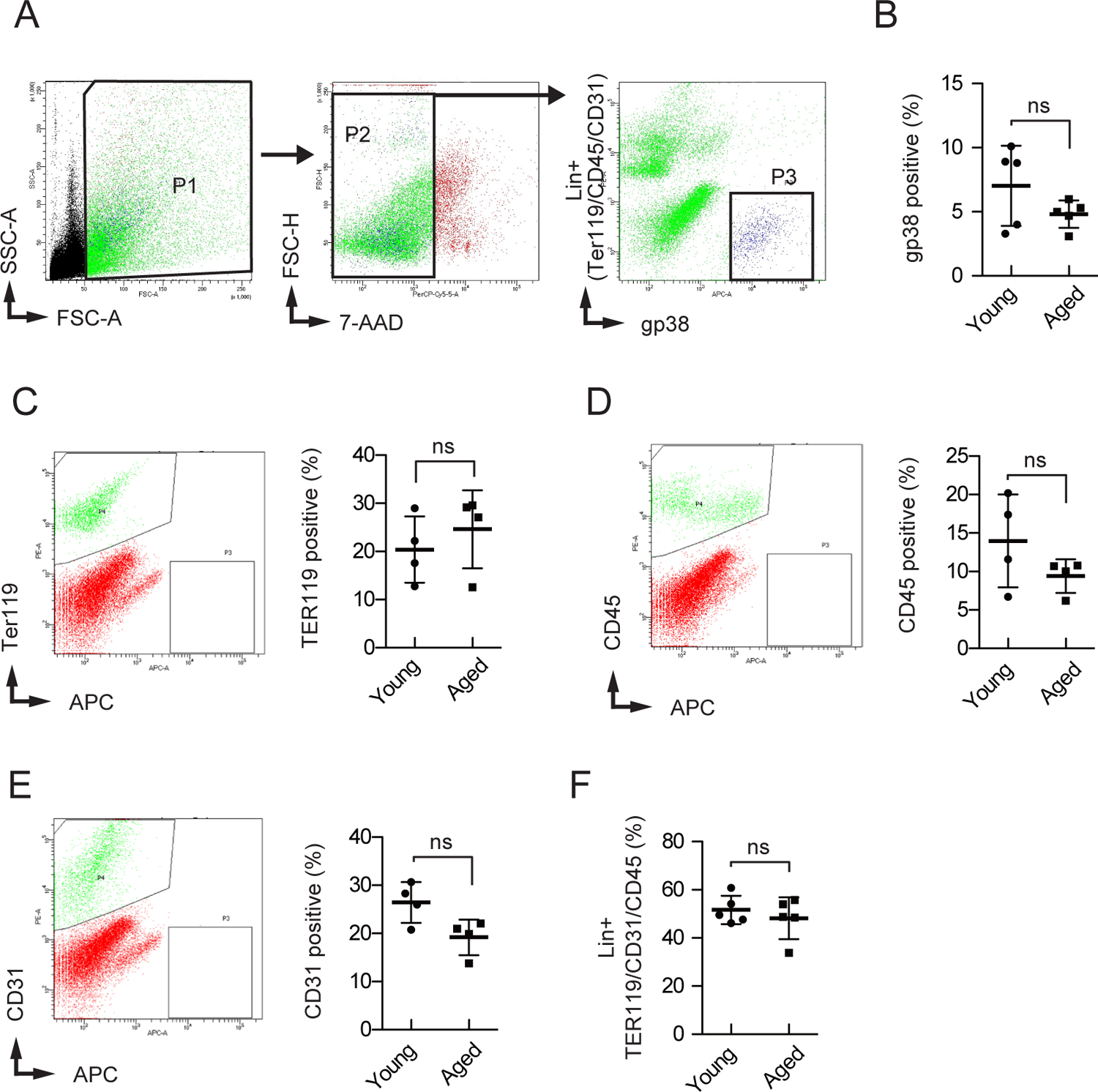
Composition of cardiac fibroblast and non-fibroblast cells in the non-CM fraction isolated from young and aged mice. A, Representative FCM analysis of cardiac fibroblasts in the non-CM fraction based on the gp38 marker. P1, the single cell fraction. P2, 7-AAD-negative live cells. P3, Lin-negative and gp38-positive cardiac fibroblasts. B, The proportion of gp38^+^ fibroblasts in the non-CM fraction remained largely consistent with aging (*n* = 5). ns, not significant. C, FCM analysis of Ter119 marker-based erythrocyte composition in young and aged non-CM fraction: representative FCM image (left) and quantitative analysis (right) (*n* = 4). D, FCM analysis of CD45 marker-based leukocyte composition in young and aged non-CM fraction: representative FCM image (left) and quantitative analysis (right) (*n* = 4). E, FCM analysis of CD31 marker-based endothelial cell composition in young and aged non-CM fraction: representative FCM image (left) and quantitative analysis (right) (*n* = 4). F, Quantitative analysis of Lin^+^ (TER119-, CD31-, and CD45-positive) non-CM cell composition in young and aged samples (*n* = 5).

### RNA sequencing revealed CM and non-CM specific changes in gene expression

We next performed transcriptome analysis of CM and non-CM fractions to elucidate specific gene expression of CM as a postmitotic cell. Because the analysis was conducted using single samples (n=1) and statistical analysis was unable, we explored candidate genes that were upregulated or downregulated using a log_2_ fold change (FC) threshold of |1| (M-value), regardless of the magnitude of mean expression levels (A-value). We found that 1104 protein-coding genes were upregulated with a log_2_FC greater than 1 in CM fraction upon aging, whereas, only 89 genes were downregulated with a log_2_FC less than −1, indicating predominant upregulation of genes (Fig. 3A). Whereas 257 protein-coding genes were upregulated with a log_2_FC greater than 1 in non-CM fraction with aging, and 844 genes were downregulated with a log_2_FC less than −1 (Fig. 3B). 70 genes were concomitantly upregulated between CM and non-CM, whereas 9 genes were downregulated. Additionally, only 39 and 15 genes exhibited counterregulation between CM and non-CM with aging (Fig. 3C).

**Figure 3.**
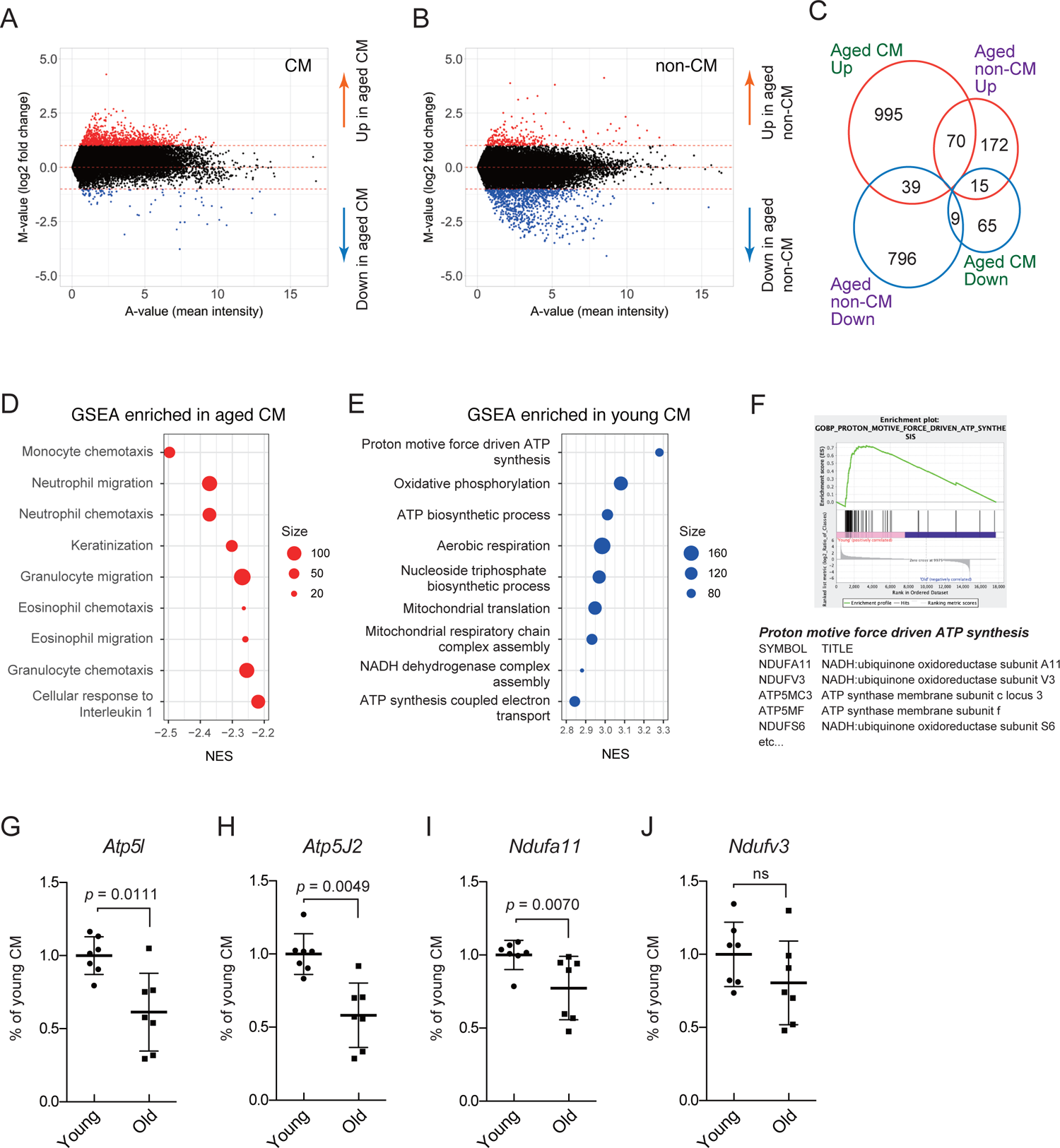
Transcriptome analysis for protein-coding genes of the CM and the non-CM isolated from young and aged mice. A, MA plot for the protein-coding genes between the young and aged CM. Upregulated (red) and downregulated (blue) genes in aged CM were defined as log2 FC > |1| (M-value). B, MA plot between the young and aged non-CM. C, Venn diagram displaying the number of upregulated and downregulated protein-coding genes in aged CM and non-CM samples, respectively. D, Summary of the GO-BP terms enriched in aged CM sample by GSEA analysis. NES, normalized enrichment score. E, Summary of the GO-BP terms enriched in young CM sample by GSEA analysis. F, GSEA plot of the top ranked GO-BP enriched in young CM sample and representative gene symbols in the gene list. G-J, qPCR analyses for two ATP synthases in mitochondrial respiratory chain complex V (*Atp5l* and *Atp5J2*) and two NADH dehydrogenases in complex I (*Ndufa11* and *Ndufv3*).

Performing GO analysis with the DAVID database on the top 1000 protein-coding genes that were either upregulated or downregulated, we observed an enrichment of inflammatory response and immune-related biological processes (BP) in the upregulated genes within the aging CM fraction (Table 1). These findings suggest the activation of inflammaging in CM. This is also the case in non-CM fraction (Table 2). On the other hand, most of downregulated genes were enriched for mitochondrial respiratory chain in CM, which is consistent with the previous report [7], suggesting a potential decline in the functional capacity of mitochondrial ATP production within CM (Table 1). Clearly different result was obtained for non-CM fraction in which downregulated genes were notably enriched in cell cycle and DNA replication process, implying occurrence of cell cycle arrest in aged non-CM (Table 2). We also employed GSEA and obtained similar results to GO-BP in which the genes were enriched in inflammatory-related process in the aged CM (Fig. 3D) and in the mitochondrial ATP biosynthetic process in the young CM sample (Figs. 3E and 3F), which indicated a reduction in respiratory activity in aged CM. To confirm downregulation of genes related to mitochondrial electron transport chain (ETC) with aging, we performed qPCR for two ATP synthases in complex V (*Atp5l* and *Atp5J2*) and two NADH dehydrogenases in complex I (*Ndufa11* and *Ndufv3*) which were identified as the candidate genes with reduced levels in aged CM by RNA-seq. Expression levels of *Atp5l*, *Atp5J2*, and *Ndufa11* in aged CM were significantly lower than those in young CM (Figs. 3G-3I). Mean expression level of *Ndufv3* was also reduced with aging (Fig. 3J), although there is no statistical significance. We also evaluated changes of expression levels of these four genes in non-CM, and found no significant reduction upon aging (Figs. S1A-S1D). In liver, *Atp5l* and *Atp5J2* were significantly reduced upon aging (Figs. S1E and S1F) whereas no changes were observed in *Ndufa11* and *Ndufv3* (Figs. S1G and S1H), suggesting that reduction of ATP synthesis in Complex V are common phenomenon between CM and hepatocyte.

**Table. 1.** Top 10 GO-BP results for up- and downregulated 1000 protein-coding genes in CM (*p*-value ranked)

| Term | % | *p*-value | Bonferroni |
| --- | --- | --- | --- |
| Upregulated (FC ranked) |  |  |  |
| GO:0006954~inflammatory response | 4.8 | 1.39E-09 | 5.56E-06 |
| GO:0002376~immune system process | 5.3 | 1.55E-07 | 6.21E-04 |
| GO:0030593~neutrophil chemotaxis | 1.7 | 4.02E-07 | 0.00161 |
| GO:0006935~chemotaxis | 2.1 | 1.76E-06 | 0.00705 |
| GO:0045944~positive regulation of transcription from RNA pol II promoter | 9.1 | 1.95E-06 | 0.00780 |
| GO:0002548~monocyte chemotaxis | 1.1 | 2.74E-06 | 0.01093 |
| GO:0045893~positive regulation of transcription, DNA-templated | 5.9 | 9.17E-06 | 0.03611 |
| GO:0044419~interspecies interaction between organisms | 0.7 | 2.37E-05 | 0.09072 |
| GO:0070098~chemokine-mediated signaling pathway | 1.2 | 2.44E-05 | 0.09323 |
| GO:0071346~cellular response to interferon-gamma | 1.7 | 3.78E-05 | 0.14060 |
| Downregulated (FC ranked) |  |  |  |
| GO:0042776~mitochondrial ATP synthesis coupled proton transport | 3.5 | 9.34E-30 | 3.42E-26 |
| GO:0009060~aerobic respiration | 2.7 | 6.39E-18 | 2.34E-14 |
| GO:0032981~mitochondrial respiratory chain complex I assembly | 2.5 | 1.40E-16 | 4.07E-13 |
| GO:0006123~mitochondrial electron transport, cytochrome c to oxygen | 1.1 | 5.96E-09 | 2.18E-05 |
| GO:0032543~mitochondrial translation | 1.9 | 1.50E-07 | 5.49E-04 |
| GO:0015986~ATP synthesis coupled proton transport | 1.0 | 2.82E-07 | 0.001 |
| GO:0046034~ATP metabolic process | 1.3 | 8.01E-07 | 0.003 |
| GO:0045333~cellular respiration | 1.0 | 9.56E-07 | 0.003 |
| GO:0006122~mitochondrial electron transport, ubiquinol to cytochrome c | 0.7 | 9.47E-06 | 0.034 |
| GO:0006120~mitochondrial electron transport, NADH to ubiquinone | 0.9 | 1.06E-05 | 0.038 |

**Table. 2.** Top 10 GO-BP results for up- and downregulated 1000 protein-coding genes in non-CM (*p*-value ranked)

| Term | % | *p*-value | Bonferroni |
| --- | --- | --- | --- |
| Upregulated (FC ranked) |  |  |  |
| GO:0006954~inflammatory response | 7.7 | 2.70E-27 | 1.07E-23 |
| GO:0002376~immune system process | 9.0 | 3.70E-27 | 1.47E-23 |
| GO:0030593~neutrophil chemotaxis | 3.1 | 2.50E-20 | 9.92E-17 |
| GO:0045087~innate immune response | 7.8 | 6.66E-13 | 2.64E-09 |
| GO:0070098~chemokine-mediated signaling pathway | 2.0 | 1.16E-12 | 4.58E-09 |
| GO:0006955~immune response | 6.2 | 3.22E-12 | 1.28E-08 |
| GO:0006935~chemotaxis | 2.9 | 5.89E-12 | 2.34E-08 |
| GO:0071347~cellular response to interleukin-1 | 2.1 | 2.56E-10 | 1.01E-06 |
| GO:0009617~response to bacterium | 4.0 | 1.12E-09 | 4.44E-06 |
| GO:0050766~positive regulation of phagocytosis | 1.9 | 2.11E-09 | 8.35E-06 |
| Downregulated (FC ranked) |  |  |  |
| GO:0007049~cell cycle | 15.1 | 2.83E-65 | 8.97E-62 |
| GO:0051301~cell division | 11.0 | 3.30E-55 | 1.05E-51 |
| GO:0006334~nucleosome assembly | 5.3 | 4.90E-40 | 1.55E-36 |
| GO:0007059~chromosome segregation | 4.1 | 5.75E-28 | 1.82E-24 |
| GO:0006335~DNA replication-dependent nucleosome assembly | 2.0 | 2.92E-21 | 9.25E-18 |
| GO:0006260~DNA replication | 3.7 | 2.21E-20 | 7.00E-17 |
| GO:0006281~DNA repair | 6.1 | 4.61E-17 | 1.46E-13 |
| GO:0006974~cellular response to DNA damage stimulus | 7.1 | 3.97E-16 | 1.41E-12 |
| GO:0007094~mitotic spindle assembly checkpoint | 1.9 | 5.23E-16 | 1.76E-12 |
| GO:0000070~mitotic sister chromatid segregation | 1.9 | 3.89E-15 | 1.23E-11 |

### Differentially expressed snoRNA genes in aged CM

We further analyzed transcriptome data to identify changes in non-coding RNAs (ncRNAs). Among several ncRNAs such as long non-coding RNA (lncRNA) and microRNA (miRNA), many snoRNAs were differentially expressed between young and aged CM (Fig. 4A). Among them, we focused on five candidate snoRNAs (Snora15, Snora24, Snora41, Snora47, and Snora62), which were highly expressed in either young or aged samples, but undetectable in the opposite age group by RNA-seq analysis. Furthermore, expression of these snoRNAs was not detected in the non-CM samples (Figs. 4B and 4C), suggesting their CM-specific expressions. We performed qPCR to evaluate expression levels of these snoRNAs and found that four snoRNAs (Snora15, Snora24, Snora41, and Snora47) were significantly downregulated in aged CM (Figs. 4D-4G), while one (Snora62) was not (Fig. 4H). None of these five snoRNAs were differentially expressed in non-CM samples with aging (Figs. 4I-4M). Snora15, which was identified as an upregulated snoRNA in aged CM sample by RNA-seq analysis (Figs. 4A and 4C), but was significantly downregulated with aging in qPCR analysis (Fig. 4G). Although the reasons are not clear, the possibility that expression level of Snora15 is highly variable among individuals cannot be ruled out.

**Figure 4.**
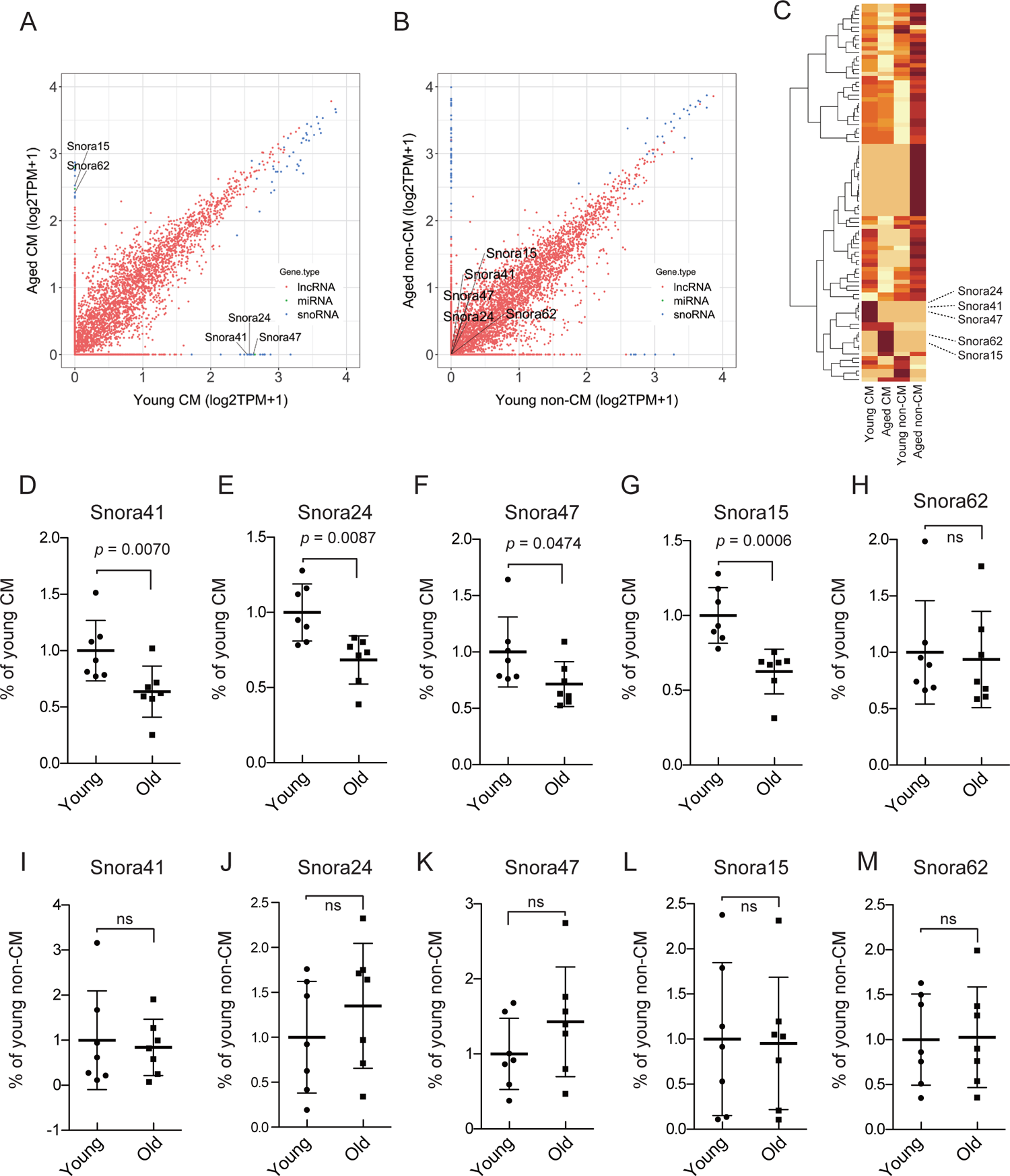
Transcriptome analysis for non-coding RNAs of the CM and the non-CM isolated from young and aged mice. A, Scatter plot for the non-coding RNAs between the young and aged CM. Candidate snoRNAs are indicated in the plot. B, Scatter plot between the young and aged non-CM. C, Heatmap analysis of differentially expressed snoRNAs in young and aged samples. The color intensity reflects the expression level, with darker colors representing higher expression and lighter colors indicating lower expression. snoRNAs that were not detected in any of the samples were excluded. D-H, qPCR analyses for candidate snoRNAs in young and aged CM. I-M, qPCR analyses for candidate snoRNAs in young and aged non-CM. ns, not significant.

## Discussion

The heart is composed of postmitotic CM and non-CM. Under physiological conditions, both cell types undergo senescence with cardiac aging, and they mutually affect senescence progression of neighboring cells by releasing distinct proinflammatory factors [10], although which cell types have the leading role in cardiac degeneration with aging and age-related diseases are still unknown. In this study, we obtained clearly distinct results by transcriptome analyses of isolated CM and non-CM in which genes involved in ATP biosynthetic process via mitochondrial ETC were significantly downregulated in CM (Figs. 3G-3J), but not in non-CM. We also identified that several snoRNAs are specifically downregulated in aged CM (Figs. 4D-4G), which could be novel senescence markers particular to postmitotic cells. It also underlines the results that compositions of fibroblasts, erythrocytes, leukocytes, and endothelial cells in the non-CM remained relatively stable with aging (Figs. 2A-2F). It is essential to note that alterations in the population of specific cell types during aging could potentially influence the observed gene expression pattern. Thus, our transcriptome analysis of the non-CM fraction appears to suggest that gene expression changes are not solely dependent on shifts in cellular composition.

Age-associated global downregulation of mitochondrial ETC and ATP synthase genes have been observed in whole ventricle [17] and isolated cardiomyocytes [7]. This possibly contributes to reduction in ETC complexes I and V activities, oxygen consumption rate, and ATP production in aged mitochondria from heart [17, 18]. Our transcriptome results support these previous findings with qPCR confirmations. Furthermore, we revealed that the expression levels of genes involved in mitochondrial complexes I and V did not significantly decrease with aging in the non-CM (Figs. S1A-S1D), while in the liver, such alterations were partly observed (Figs. S1E-S1F). Therefore, the decline in mitochondrial function observed in the cardiac aging appears to be specific to CM, and it is strongly suggested that non-CM are not directly involved in age-related cardiac mitochondrial dysfunction, although paracrine effects of SASP from non-CM to CM are not negligible. On the other hand, in the liver, several mitochondrial genes were reduced with aging. Differential expression of genes encoding mitochondrial ETC subunits [19] and decrease in activities of some ETC complexes [20] in various aged tissues including the liver have been reported. Further elucidation will be necessary to determine whether these changes in gene expression associated with aging are due to a common mechanism among tissues, and whether this leads to establish an optimal balance between aerobic ATP synthesis and the suppression of excess ROS production from mitochondria.

snoRNAs are deeply involved in RNA modification reactions such as pseudouridylation stabilizing rRNA and mRNA splicing, and have been eminently studied from the perspective of its utility as a tumor marker [21–24]. Due to the critical role of snoRNAs in protein synthesis on ribosome, their proper expression significantly influences cellular proliferation and differentiation [25, 26]. Our study demonstrated that some snoRNAs could be one of novel aging markers in CM or postmitotic cells. By identifying interacting protein factors and other components, it is expected that the individual functions of the identified snoRNAs and their roles in aging will be uncovered in the future.

In conclusion, there is a significant gap between the gene profiles of CM and non-CM with aging, suggesting distinct aging processes exist. Extending this idea, the variances between different tissues are notably pronounced, suggesting the possible existence of diverse senescence mechanisms unique to each tissue. Given these observations, defining general concept of senescence in postmitotic cells, especially in differentiated cells, becomes highly challenging. Nonetheless, we try to define senescence of postmitotic cells as the irreversible decline in the specialized functions in differentiated cells.

## Acknowledgement

The authors are grateful to Santa Cruz Biotechnology, Inc. (Dallas, TX) and Cosmo Bio Co., Ltd. (Tokyo, Japan) for providing the sample antibodies listed in Supporting Information. The authors would like to thank Ms. Masumi Shimizu for aiding FCM analyses, Dr. Mika Sakamoto of National Institute of Genetics for aiding RNA-seq data analyses, and Editage (www.editage.jp) for English language editing. Computations were partially performed on the NIG supercomputer at ROIS National Institute of Genetics. We would like to express my sincere gratitude to ChatGPT (GPT-4), developed by OpenAI, for its invaluable assistance in editing the English text.

## Author contributions

YT and YK designed the research and drafted the paper; YT carried out all experiments and analyzed data; MH analyzed transcriptome data; HO, II, and MI supported the cell culture and animal experiments.

## Supporting information

### 1. Quantitative PCR (qPCR) primers

Mouse *Tnnc1*

Mm_Tnnc1_RT-UP1, 5’-ATGATTGACGAAGTAGACGAGGA-3’

Mm_Tnnc1_RT-LP1, 5’-TCATAGTCAATTCGGCCATCGTT-3’

Mouse *αMHC*

Mm_Myh6(aMHC)_RT-UP1, 5’-GCCGTATCATCACCAGAATCCAG-3’

Mm_Myh6(aMHC)_RT-LP1, 5’-GCATCTTTGACTCGCCCAA-3’

Mouse *Col1a1*

Mm_Col1a1(collagen1)_RT-UP1, 5’-GCATAAAGGGTCATCGTGGCTTC-3’

Mm_Col1a1(collagen1)_RT-LP1, 5’-GCTGTCACCAGTGCGACCTC-3’

Mouse *αSMA*

Mm_Acta2(aSMA)_RT-UP1, 5’-ACCATGTACCCAGGCATTGCTGA-3’

Mm_Acta2(aSMA)_RT-LP1, 5’-ATCCAGACAGAGTACTTGCGTTC-3’

Mouse *Atp5l*

Mm_Atp5l_RT-UP1, 5’-CGTACCTTCCACCTTAGACCA-3’

Mm_Atp5l_RT-LP1, 5’-CCAGCTCAACCTTGGCGTA-3’

Mouse *Atp5J2*

Mm_Atp5j2_RT-UP1, 5’-CTTCAAGATGGCGTCACTCGT-3’

Mm_Atp5j2_RT-LP1, 5’-GGACCATGCTAATCCCCGAGA-3’

Mouse Ndufa11

Mm_Ndufa11_RT-UP1, 5’-GACCTACATCACCACGGCCTT-3’

Mm_Ndufa11_RT-LP1, 5’-GCCAAACATCGCTCCAATAGCAG-3’

Mouse *Ndufv3*

Mm_Ndufv3_RT-UP1, 5’-CTACAGAGTCAGAGAAGAGTGCAA-3’

Mm_Ndufv3_RT-LP1, 5’-CTGTGTTCTCCACTGATGGGT-3’

Mouse Snora41

Mm_Snora41_RT-UP1, 5’-GCTGTCTTTATGGTAGCAGT-3’

Mm_Snora41_RT-LP1, 5’-AGGCATTAAATTGTGTAACAGT-3’

Mouse Snora24

Mm_Snora24_RT-UP1, 5’-GGCAGTCTCCCTTCCTAGCCATG-3’

Mm_Snora24_RT-LP1, 5’-ATGAAATGGTGACAGCTTTGCCAA-3’

Mouse Snora47

Mm_Snora47_RT-UP1, 5’-TGCTGCCTTCCATTGGTTAAGACCT-3’

Mm_Snora47_RT-LP1, 5’-TCAGTGCTGCCCCTCCACGG-3’

Mouse Snora15

Mm_Snora15_RT-UP1, 5’-TAATTTACACAGCCAGACACC-3’

Mm_Snora15_RT-LP1, 5’-CCATAAAATACAGCATATCAGCC-3’

Mouse Snora62

Mm_Snora62_RT-UP1, 5’-CGCTGGTCGATGAACTCACA-3’

Mm_Snora62_RT-LP1, 5’-GTTACTATCAAACCCCACTCACT-3’

For 18S ribosomal RNA

Univ-18S-F, 5’-AGCTTGCGTTGATTAAGTCCCT-3’

Univ-18S-R, 5’-GCCTCACTAAACCATCCAATCGG-3’

---

**Figure S1.**
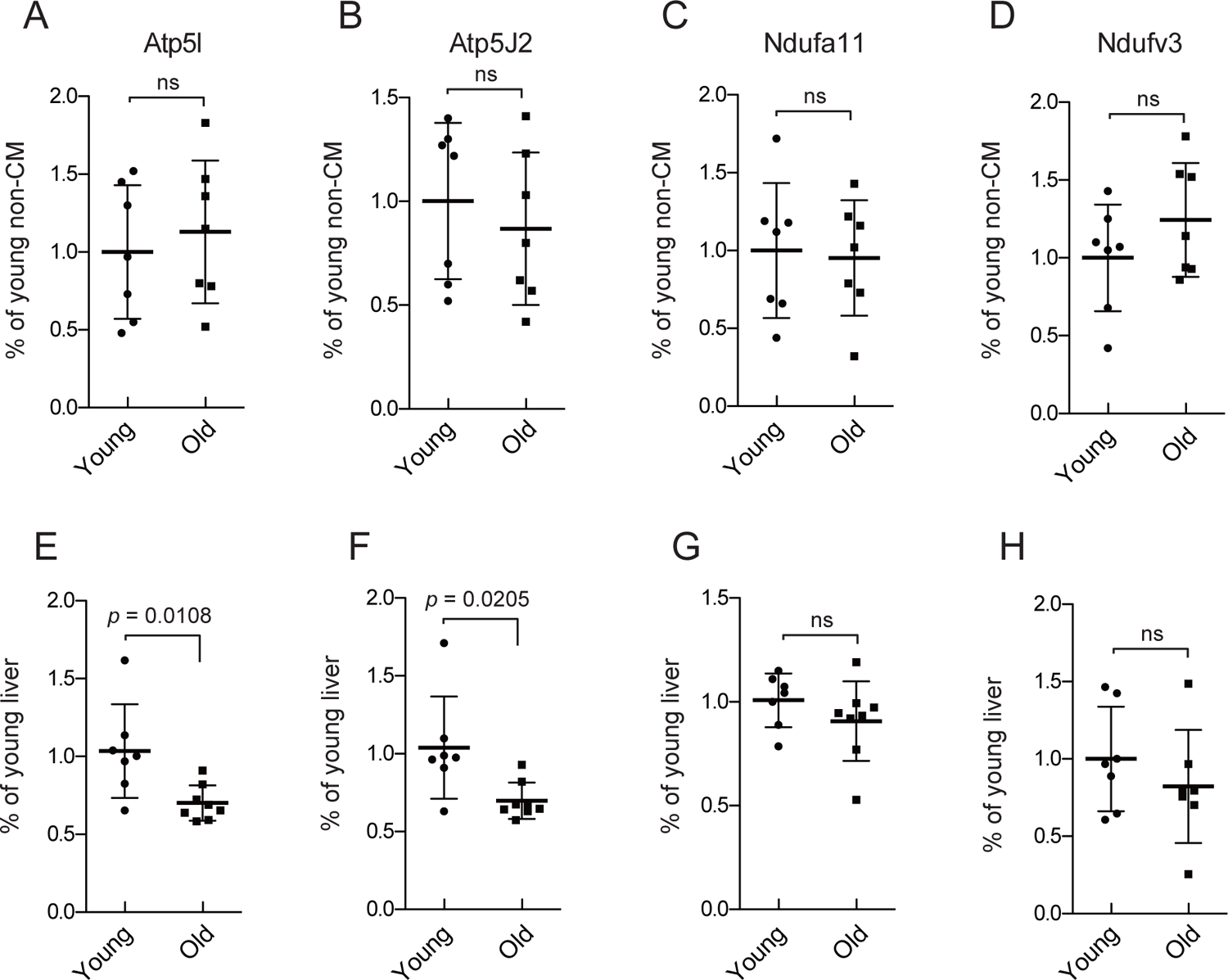

